# ANTIMICROBIAL ACTIVITY OF CLOVE EXTRACTS AGAINST MICROORGANISMS ISOLATED FROM VAGINAL DISCHARGE

**DOI:** 10.1101/2024.06.13.598938

**Authors:** Aishat Taiye Omoniyi, Muneer Yaqub

## Abstract

Infections of the genitourinary and reproductive tracts pose significant health concerns for women, particularly those of reproductive age. These infections often manifest as vaginal discharge and can be caused by a variety of microorganisms, including pathogenic bacteria and fungi. Traditional antibiotic treatments are increasingly challenged by the rise of antibiotic-resistant strains, underscoring the need for alternative therapies. This study aimed to isolate and identify microorganisms from vaginal swab samples and evaluate the antimicrobial efficacy of clove (*Eugenia caryophyllata*) extracts against these isolates. Using CLED agar, nutrient agar, and Sabouraud dextrose agar, a diverse range of bacterial and fungal flora were isolated from eight vaginal swab samples. The primary bacterial isolates included Proteus mirabilis, Escherichia coli, Staphylococcus aureus, and Lactobacillus spp., while Candida albicans was the main fungal isolate. Biochemical tests confirmed the identity of these microorganisms. The study found that ethanol clove extract exhibited significant antimicrobial activity, particularly against Staphylococcus aureus and Candida albicans, with the minimum inhibitory concentration (MIC) values being 20 mg/ml and 10 mg/ml, respectively. Additionally, ciprofloxacin, used as a control antibiotic, showed maximum inhibition against Lactobacillus spp., highlighting a potential risk for disrupting beneficial vaginal flora when using conventional antibiotics. The findings suggest that ethanol clove extract could serve as an effective alternative antimicrobial agent, reducing the risk of antibiotic resistance and preserving the balance of the vaginal microbiome. This research emphasizes the importance of exploring plant-based antimicrobial agents as viable alternatives to traditional antibiotics. The significant antimicrobial properties of clove extract against common vaginal pathogens offer promising implications for future therapeutic applications in managing vaginal infections.

## INTRODUCTION

Infections of the genitourinary and reproductive tracts are significant concerns for women’s sexual health. These infections commonly occur in women of reproductive age and often manifest as vaginal discharge [1]. Vaginal discharge in this age group is the most frequent complaint encountered by gynecologists and general practitioners [2]. Despite the control exerted by lactobacilli in the vaginal microenvironment, various other microorganisms can be cultured from healthy women’s vaginal samples [3]. While these may not always trigger a pathological state, an imbalance resulting from the dominance of one class of these organisms can lead to vaginosis or vaginitis [4].

Infections of the genitourinary or reproductive tract that can lead to vaginal discharge include sexually transmitted infections (STIs), bacterial vaginosis (BV), aerobic vaginitis, and candidiasis [5]. STIs occur when sexually transmitted organisms are introduced into the vagina, primarily through sexual activity [6,7]. This is a major issue for women of reproductive age worldwide, especially in developing countries in Africa [8]. According to WHO estimates, 75% to 80% of all new cases of sexually transmitted diseases occur in developing countries [9].

Contrary to the general knowledge of antibiotics belonging to quinolones, aminoglycosides, cephalosporins, and beta-lactams being used as treatment therapy [10], genital tract infections are challenging to resolve due to the frequent use of antibiotics [11], which ultimately leads to the development of drug resistance in pathogenic bacteria. The development of resistance by these pathogenic bacteria usually occurs via three major mechanisms: horizontal gene transfer (plasmids, transposons, and bacteriophages), incorporation of foreign DNA into bacterial DNA, and the formation of a new combination of genomic material and changes in nucleotide sequences in chromosomes, known as mutation [12]. Antibiotic resistance in these pathogens and the complications it causes in the biological system emphasize the need for alternative therapies.

Nowadays, more than 1,350 plants with antimicrobial activities and over 30,000 antimicrobial components have been extracted from plants [13]. Plants’ secondary metabolites contain numerous antimicrobial agents, making them effective against both Gram-positive and Gram-negative bacteria [14]. Particular attention should be paid to the plant kingdom, which offers many compounds with antibacterial effects that have been proven effective in treating bacterial infections without the adverse effects specific to conventional antibiotics [15]. Herbal treatments offer new chemotherapeutic alternatives due to their ability to minimize adverse drug effects, potential antimicrobial properties, and reduction of infections caused by drug-resistant pathogenic microorganisms [16].

*Eugenia caryophyllata*, commonly known as clove, is the largest genus of the Myrtaceae family. It is a tropical and subtropical flowering plant widely distributed in Asia, Africa, Madagascar, and the Pacific and Oceanic regions [17]. Clove, a precious spice derived from this plant, contains phenylpropene eugenol, which gives clove its strong characteristic aroma. This component also demonstrates broad antimicrobial activity against Gram-positive, Gram-negative, and acid-fast bacteria, as well as antifungal properties [18]. Flavonoids in clove also contribute to various cellular defense mechanisms, acting against different inflammatory mediators and free radical species [19].

The advancement of native medicines and the use of medicinal plants carry significant potential in the treatment of various ailments [20]. This is particularly relevant due to the emergence and re-emergence of microbial infections that demonstrate resistance to various antibiotics, making the exploration of new antimicrobial sources necessary. The aim of this research was to isolate and identify microorganisms causing abnormal vaginal discharge and determine the antimicrobial effect of clove extracts on these organisms.

## MATERIALS AND METHODS

### Samples Collection and clove bud processing

Vaginal swab samples were aseptically collected from women of reproductive age residing in the Usmanu Danfodiyo University female hall of residence. These samples were carefully placed into sterile swab stick containers and transported to the microbiology laboratory at Usmanu Danfodiyo University, Sokoto, Nigeria, for analysis. Cloves were procured from Sokoto Central Market, Sokoto South Local Government Area, Sokoto State, Nigeria [21]. The authenticity of the purchased cloves was verified in the Botany unit of the Biology Department at Usmanu Danfodiyo University. The dried cloves were thoroughly washed with distilled water to eliminate dust and surface contaminants, dried adequately, and pulverized into a fine powder using a mortar and pestle. The powder was then sieved and carefully packaged in an airtight container, duly labeled. The dried clove buds were ground to powder using a mortar and pestle and weighed. The powdered sample was then stored in an airtight container for further use.

### Extraction Process

The two extracts, aqueous and ethanol, were obtained from dried clove buds. For aqueous extract, the well-air-dried clove buds were ground, and 50g of the powdered clove was suspended in 300ml sterilized distilled water in a conical flask, corked with cotton wool and foil paper, and left for 3 days with regular shaking intervals. The mixture was then filtered using Whatman filter paper, and the crude aqueous extract was obtained after evaporation to dryness using a water bath. The extracts were stored at 4°C in a freezer until needed for further experiments. For the ethanol extract, 50g of the powdered clove was soaked in 300ml of ethanol and allowed to stand for 24 hours with occasional stirring. The mixture was filtered using a muslin cloth, and the filtrate was dried in a hot-air oven at 45°C [23]. Concentrated extracts of both aqueous and ethanol extract initially obtained and kept in sterile amber bottles were suspended in their respective plain agar and diluted into different concentrations (80mg, 40mg, and 20mg) to test for the antimicrobial activity of each extract.

### Isolation and Identification of Microorganisms

A sterile wire loop was used to transfer a small portion of the vaginal swab samples onto the surface of the agar in a zigzag or streak pattern. The inoculated plates were labeled for identification and incubated at 37°C for 24 hours for bacterial culture and at 25°C - 27°C for 3 days for fungal culture. Using a sterile inoculating wire loop, colonies from the overnight culture of the test organisms were transferred into a tube containing about 5ml of sterile distilled water. The turbidity was adjusted to achieve standard turbidity equal to that of the 0.5 McFarland standard. Identification was performed using Gram staining [24], biochemical tests [25], and sugar fermentation tests [26] to determine the types and quantities of microorganisms present [27].

### Determination of Antibacterial Activity

The antibacterial activity of the clove extracts was evaluated using the agar well diffusion technique. Mueller-Hinton agar plates were prepared and solidified. Bacterial suspensions were adjusted to the appropriate turbidity and inoculated onto the agar surface using a sterile swab. After inoculation, the plates were left to stand for 15 minutes to allow absorption. Three wells, each 6 mm in diameter, were bored into the agar using a sterile cork borer. Various concentrations of the ethanolic clove extract (80mg/ml, 40mg/ml, and 20mg/ml) were added to the wells. A ciprofloxacin disc was used as a positive control. The Petri dishes were incubated at 37°C overnight. The antibacterial efficacy of the clove extracts was determined by measuring the diameters of the inhibition zones in millimeters. The same procedure was repeated for aqueous concentrations of the extract [28, 29].

### Determination of Antifungal Activity

Antifungal activity was assessed using the agar well diffusion method. Sabouraud dextrose agar was poured into Petri dishes and allowed to solidify. A sterile cork borer was used to create wells in each agar plate. Fungal cultures were evenly distributed over the agar surface using a sterile swab. The plates were left to dry for 15 minutes before testing. Various concentrations of the clove extract were introduced into the wells. The plates were labeled and incubated overnight at 37°C. The antifungal efficacy of the extracts was evaluated by measuring the diameters of the inhibition zones in millimeters [30].

### Minimum Inhibitory Concentration (MIC)

The MIC of the clove extract was determined using the procedure outlined by the National Committee for Clinical Laboratory Standards. The experiment utilized twelve test tubes, each containing 5ml of peptone water. Different concentrations of the ethanolic clove extract were prepared and distributed among the tubes as follows: 80mg/ml, 40mg/ml, 20mg/ml, 10mg/ml, 5mg/ml, 2.5mg/ml, 1.25mg/ml, 0.6125mg/ml, and 0.312mg/ml. Each concentration was mixed thoroughly in its respective tube. To assess the antibacterial activity, 0.1ml of broth cultures of the test organism were added to nine of these tubes (tubes 2 to 10). Tube 1, serving as the positive control, contained broth, antibiotic, and culture. Tube 11 served as the sterility control, containing only broth, while tube 12 served as the negative control, containing broth and culture but no extract. The tubes were incubated overnight at 37°C. Post incubation, the tubes were observed for bacterial growth. The MIC was defined as the lowest concentration of clove extract that completely inhibited the visible growth of the bacteria in the inoculated tubes [31].

## RESULTS

### Isolation and Identification from Vaginal Swab Samples using CLED Agar and Nutrient Agar

Bacteria isolated from vaginal swab samples using CLED agar and nutrient agar were identified based on Gram reaction, classifying them as either Gram-positive or Gram-negative. Further identification was achieved through biochemical tests, as detailed in **Table 1**. Out of eight samples collected from females of reproductive age residing in a hostel, seven exhibited microbial growth after 24 hours of incubation.

**Table 1:** Identification of Bacteria Isolates. This table shows the identification of bacterial isolates from vaginal swab samples. The isolates were classified based on Gram reaction, morphology, and various biochemical tests, including MR (Methyl-red), VP (Voges-Proskauer), catalase, citrate, urease, sucrose, lactose, glucose, motility, H2S (Hydrogen Sulphide) production, gas production, indole, coagulase, and starch hydrolysis.

| Sample | Gram | Morphology | MR | VP | Catalase | Citrate | Urease | Sucrose | Lactose | Glucose | Motility | H<sub>2</sub>S | Gas | Indole | Coagulase | Starch | Isolated organism |
| --- | --- | --- | --- | --- | --- | --- | --- | --- | --- | --- | --- | --- | --- | --- | --- | --- | --- |
| A<sub>I</sub> | - | R o d | - | + | + | - | + | - | - | + | + | + | - | - | - | - | *Proteus mirabilis* |
| B<sub>C</sub> | - | R o d | + | - | + | - | - | + | - | - | + | - | + | + | - | - | *Escherichia Coli* |
| C<sub>B</sub> | + | Cocci | - | + | + | - | + | + | + | + | - | - | + | - | + | + | *Staphylococcus aureus* |
| D<sub>F</sub> | + | R o d | - | - | - | - | - | + | - | + | - | - | - | - | - | - | *Lactobacillus spp* |

Keys: Positive (+); Negative (-)

The study revealed a diverse microbial flora. Proteus mirabilis, a Gram-negative, rod-shaped bacterium, was identified in Sample AI. It tested negative for methyl-red (MR), positive for Voges-Proskauer (VP), catalase, urease, glucose, and motility, aligning with characteristics of an opportunistic pathogen often found in the vaginal flora. Escherichia coli, another Gram-negative, rod-shaped bacterium, was identified in Sample BC. This isolate tested positive for MR, catalase, sucrose, glucose, gas, and indole production. These results are typical of E. coli, a common commensal of the gastrointestinal tract that can also be found in the vaginal flora. Staphylococcus aureus, a Gram-positive, cocci-shaped bacterium, was identified in Sample CB. It tested positive for VP, catalase, citrate, urease, sucrose, lactose, glucose, indole, and coagulase. S. aureus is a common skin and mucous membrane commensal that can become pathogenic. Lactobacillus spp., a Gram-positive, rod-shaped bacterium, was identified in Sample DF. This isolate tested positive for sucrose and glucose while being negative for other tests. Lactobacillus species are predominant in the vaginal flora, playing a crucial role in maintaining vaginal health by producing lactic acid and maintaining an acidic pH.

The presence of commensal bacteria like Lactobacillus spp. is essential for a healthy vaginal microbiome. However, the isolation of potential pathogens such as Proteus mirabilis, Escherichia coli, and Staphylococcus aureus indicates a delicate balance within the vaginal flora. These bacteria can become pathogenic under certain conditions, suggesting a potential risk of infections, especially in immunocompromised individuals or those with disrupted vaginal microbiota.

### Frequency of Occurrence of Bacterial Isolates in Vaginal Swabs

To understand the prevalence of different bacterial isolates identified in vaginal swab samples, the frequency and potential implications for vaginal health were analyzed. As shown in **Table 2**, the most common Gram-negative bacterium isolated was Proteus mirabilis, accounting for 33.33% of the isolates, followed by Escherichia coli at 16.66%. Among the Gram-positive bacteria identified, Staphylococcus aureus was the most common, also at 33.33%, followed by Lactobacillus spp. at 16.66%.

**Table 2:**
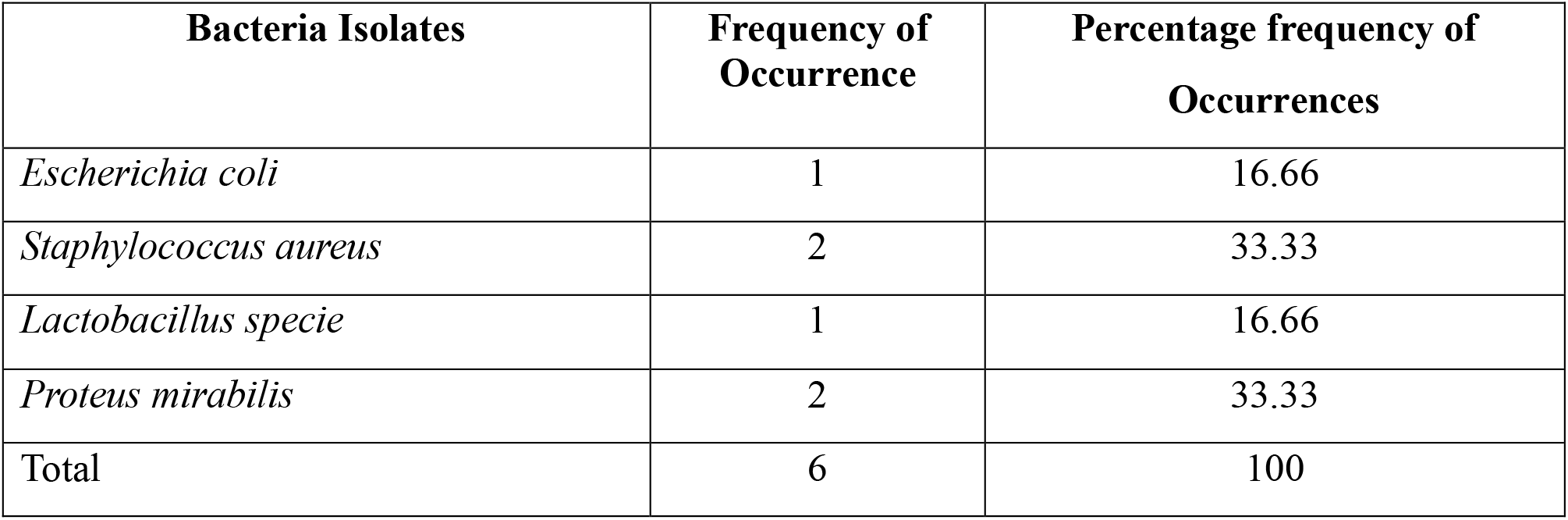
Frequency and Prevalence of Bacterial Isolates. This table presents the frequency and percentage occurrence of bacterial isolates identified from vaginal swab samples.

The presence of Lactobacillus spp. in one of the samples indicates that not all samples revealed the growth of pathogenic microorganisms. This observation is crucial as it highlights that not all cases of vaginal discharge reported by women during clinical visits are due to disease conditions or caused by pathogenic organisms [31]. The prevalence of Proteus mirabilis aligns with previous findings by Verner et al., [32] where Proteus mirabilis showed the highest frequency of occurrence among bacterial isolates.

The distribution of bacterial isolates emphasizes the microbial diversity within the vaginal flora. While pathogenic bacteria like Proteus mirabilis and Staphylococcus aureus are present, the detection of Lactobacillus spp. underscores the role of beneficial bacteria in maintaining vaginal health. This balance is essential for preventing infections and promoting a healthy vaginal environment.

### Isolation and Identification of Fungi in Vaginal Swabs: Morphological and Biochemical Characteristics

Building on the bacterial identification, this experiment also aimed to identify fungal isolates to provide a comprehensive understanding of the microbial environment in vaginal swabs. The macroscopic, microscopic, and biochemical identification (sugar fermentation) of the fungal isolate cultured on Sabouraud dextrose agar is shown in **Table 3**. Based on colony morphology,

**Table 3:** Identification of Fungal Isolate. This table details the identification of the fungal isolate from vaginal swab samples based on macroscopic, microscopic, and biochemical characteristics (sugar fermentation). The fungal isolate identified was Candida albicans, characterized by a cream-colored, smooth colony and the presence of a spherical budding yeast-like cell. The table includes results for the germ tube test and sugar fermentation of glucose, sucrose, galactose, maltose, and lactose.

| Sample | Macroscopy | Microscopy | Germ tube | Sugar fermentation |  |  |  |  | Isolate identified |
| --- | --- | --- | --- | --- | --- | --- | --- | --- | --- |
|  |  |  |  | Glucose | Sucrose | Galactose | Maltose | Lactose |  |
| C<sub>B</sub> | Cream coloured and smooth colony | Presence of a spherical budding yeast like cell | + | + | - | + | + | - | *Candida albican* |

**Keys:** Positive (+); Negative (-)

Gram staining, germ tube test, and sugar fermentation tests, the results indicated the presence of Candida albicans. Candida albicans was isolated from one of the eight samples. Within 72 hours of incubation, there was rapid growth of the isolate, displaying a cream-colored, pasty, and smooth appearance on macroscopic examination.

These findings indicate that vaginal swab cultures in this study yielded mainly aerobic bacteria and fungi, aligning with previous studies on the microbial and antibiotic sensitivity patterns of vaginal swab cultures [33]. The aerobic bacteria are typically seen in cases of aerobic vaginitis. However, no anaerobic organisms, which are usually associated with bacterial vaginosis, were detected.

While both aerobic vaginitis and bacterial vaginosis are associated with vaginal discharge, aerobic vaginitis is marked by clinical signs of inflammation, including yellowish discharge and vaginal dyspareunia. In both conditions, there is a depletion of Lactobacillus in the vagina [34]. Lactobacillus, a normal flora, serves as a check for other pathogenic microorganisms and can be displaced by activities such as sexual activity and indiscriminate use of antibiotics, thereby exposing the vagina to infections [34].

### Antimicrobial Activity of Clove Extracts on Isolated Microorganisms from Vaginal Swabs

Plant extracts can serve as an alternative and sustainable solution to control the growth of pathogenic microorganisms [35]. The results from this study, as shown in **Table 4**, indicated that ethanol clove extract has significant antimicrobial activity on some of the isolates (Escherichia coli, Staphylococcus aureus, and Candida albicans) at a high concentration of 80 mg/ml. The maximum zone of inhibition (18 mm) was observed against S. aureus at 80 mg/ml, while the minimum zone of inhibition (7 mm) was against Candida albicans at a 40 mg/ml concentration, with no visible inhibitory effect on Lactobacillus spp. and Proteus mirabilis.

**Table 4:**
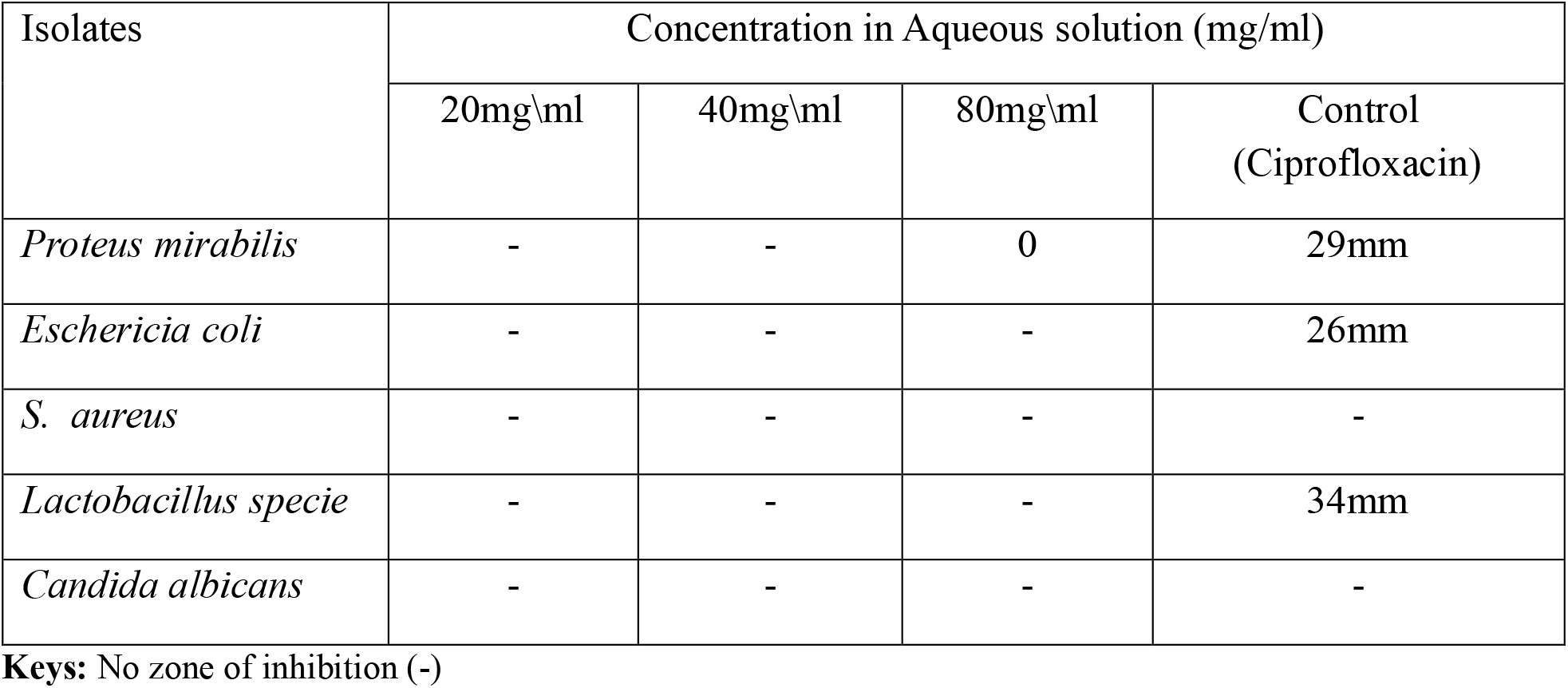
Diameter of Inhibition Zone (Aqueous Clove Extract) on the Isolates. This table illustrates the antimicrobial activity of aqueous clove extract against various bacterial and fungal isolates from vaginal swab samples. The table shows the diameter of the inhibition zones for Proteus mirabilis, Escherichia coli, Staphylococcus aureus, Lactobacillus spp., and Candida albicans at different concentrations of the aqueous clove extract (20 mg/ml, 40 mg/ml, 80 mg/ml) and a control (ciprofloxacin).

The aqueous clove extract showed less activity against all isolates, which may be due to the in-vitro assay conditions of the clove extract [36]. This contrasts with previous studies where both aqueous and ethanol clove extracts were shown to inhibit the growth of pathogenic microorganisms [37]. These results suggest that ethanol clove extract could be a potent antimicrobial agent against certain vaginal pathogens.

### Antibiotic Susceptibility of Vaginal Isolates and Implications for Flora Depletion

The antibiotic ciprofloxacin, used as a control in the antimicrobial activity testing of clove extracts, demonstrated significant inhibitory effects on several isolates. Notably, ciprofloxacin showed maximum inhibition on Lactobacillus spp. with a 34 mm zone of inhibition **(Table 5)**. This is in contrast with most studies, which have shown that Lactobacillus is generally resistant to ciprofloxacin [38, 39]. This discrepancy may be attributed to the specific species of Lactobacillus isolated in this study, suggesting variability in antibiotic susceptibility among different Lactobacillus species.

**Table 5:** Diameter of Inhibition Zone (Ethanol Clove Extract) on the Isolates. This table shows the antimicrobial activity of ethanol clove extract against various bacterial and fungal isolates from vaginal swab samples. The table lists the diameter of the inhibition zones for Proteus mirabilis, Escherichia coli, Staphylococcus aureus, Lactobacillus spp., and Candida albicans at different concentrations of the ethanol clove extract (20 mg/ml, 40 mg/ml, 80 mg/ml) and a control (ciprofloxacin). The results highlight the significant inhibition of Staphylococcus aureus and Candida albicans by the ethanol clove extract.

| Isolates | Concentration in Ethanol solution (mg/ml) |  |  |  |
| --- | --- | --- | --- | --- |
|  | 20mg\ml | 40mg\ml | 80mg\ml | Control<br>(Ciprofloxacin) |
| *Proteus mirabilis* | - | - | - | 29mm |
| *Eschericia coli* | - | 11mm | 14mm | 26mm |
| *S. aureus* | - | 16mm | 18mm | - |
| *Lactobacillus specie* | - | - | - | 34mm |
| *Candida albicans* | - | 7mm | 10mm | - |

**Keys:** No zone of inhibition (-)

The results from Table 5 highlight the potent activity of ciprofloxacin against not only pathogenic bacteria but also beneficial microorganisms such as Lactobacillus spp. The indiscriminate use of ciprofloxacin could therefore lead to the depletion of the beneficial vaginal flora, which plays a critical role in maintaining vaginal health by producing lactic acid and maintaining an acidic pH that inhibits the growth of pathogenic bacteria.

This depletion of Lactobacillus spp. could have significant implications for vaginal health. Lactobacillus spp. are essential for a healthy vaginal microbiome, and their reduction can lead to an increased risk of infections by opportunistic pathogens [40] such as Proteus mirabilis, Escherichia coli, Staphylococcus aureus, and Candida albicans, all of which were identified in the vaginal swab samples from this study.

Moreover, the study demonstrated that ethanol clove extract had significant antimicrobial activity against Escherichia coli, Staphylococcus aureus, and Candida albicans at a high concentration of 80 mg/ml, suggesting that plant extracts could serve as a viable alternative to conventional antibiotics like ciprofloxacin. The use of such natural products could help mitigate the adverse effects of antibiotics on beneficial microorganisms and reduce the risk of disrupting the vaginal or microbiome.

In summary, while ciprofloxacin is effective against various pathogenic microorganisms, its impact on beneficial bacteria like Lactobacillus spp. underscores the need for cautious and judicious use of antibiotics. Exploring alternative treatments such as clove extracts could offer a more sustainable approach to managing vaginal infections while preserving the delicate balance of the vaginal flora.

### Minimum Inhibitory Concentration (MIC) of Ethanol Clove Extract against Vaginal Isolates

To further explore the effectiveness of ethanol clove extract against vaginal isolates, the Minimum Inhibitory Concentration (MIC) was determined. The MIC is recorded as the lowest concentration of the extract that prevents visible growth of the inoculum, as shown in **Table 6**. Various concentrations were tested, with 80 mg/ml being the highest and 0.312 mg/ml being the lowest.

**Table 6:** Minimum Inhibitory Concentration of Ethanol Clove Extract. This table displays the Minimum Inhibitory Concentration (MIC) of ethanol clove extract against various vaginal isolates. The table indicates the lowest concentration of the extract that prevents visible growth of the microorganisms. The tested concentrations range from 80 mg/ml to 0.312 mg/ml. The results show that Candida albicans had the lowest MIC value at 10 mg/ml, while Staphylococcus aureus and Escherichia coli both had MIC values of 20 mg/ml. The table highlights the effectiveness of ethanol clove extract in inhibiting the growth of these common vaginal pathogens.

| Isolates | Concentration of CEE used mg\ml |  |  |  |  |  |  |  |  |  |
| --- | --- | --- | --- | --- | --- | --- | --- | --- | --- | --- |
|  | 80 | 40 | 20 | 10 | 5 | 2.5 | 1.2 | 0.1 | 0.3 | MIC |
| *Staphylococcus aureus* | - | - | - | + | + | + | + | + | + | 20mg\ml |
| *Escherichia coli* | - | - | - | + | + | + | + | + | + | 20mg\ml |
| *Candida albicans* | - | - | - | - | + | + | + | + | + | 10mg\ml |

**Keys:** Clove ethanol extract (CEE); Minimum inhibitory concentration (MIC); No growth or turbidity (-); Presence of growth or turbidity (+)

The results indicated that the lowest MIC value recorded for ethanol clove extract was 10 mg/ml for Candida albicans, while the highest MIC value was 20 mg/ml for both Staphylococcus aureus and Escherichia coli. These findings suggest that ethanol clove extract is particularly effective against Candida albicans at lower concentrations compared to other isolates. These findings reinforce the potential of ethanol clove extract as an effective antimicrobial agent against certain vaginal pathogens, with varying degrees of effectiveness depending on the microorganism.

## DISCUSSION

This study aimed to isolate and identify microorganisms from vaginal swab samples and assess the antimicrobial activity of clove extracts against these isolates. Infections of the genitourinary and reproductive tracts are a significant concern for women’s health, particularly in those of reproductive age, where vaginal discharge is a common complaint. Despite the protective role of lactobacilli, various microorganisms can inhabit the vaginal environment, and an imbalance can lead to conditions such as vaginosis or vaginitis.

The study identified a diverse range of bacterial and fungal flora from vaginal swab samples using CLED agar, nutrient agar, and Sabouraud dextrose agar. Proteus mirabilis, Escherichia coli, Staphylococcus aureus, and Lactobacillus spp. were isolated from the samples. The identification was based on Gram reaction, colony morphology, and biochemical tests. The presence of Lactobacillus spp. in some samples underscores their role in maintaining vaginal health by producing lactic acid and maintaining an acidic pH. However, pathogenic bacteria such as Proteus mirabilis, Escherichia coli, and Staphylococcus aureus were also present, indicating a potential risk for infections, especially when the vaginal microbiome is disrupted.

The study revealed that Proteus mirabilis and Staphylococcus aureus were the most frequently isolated bacteria, each accounting for 33.33% of the isolates. Escherichia coli and Lactobacillus spp. were less frequent, each at 16.66%. These findings highlight the microbial diversity within the vaginal environment and emphasize the need for careful monitoring and management of vaginal health to prevent infections.

The ethanol clove extract demonstrated significant antimicrobial activity against Escherichia coli, Staphylococcus aureus, and Candida albicans at a high concentration of 80 mg/ml, with the maximum zone of inhibition observed against Staphylococcus aureus (18 mm). The aqueous clove extract showed less activity, which may be attributed to the extraction method and assay conditions. These results suggest that ethanol clove extract could be an effective alternative to conventional antibiotics for managing vaginal infections, particularly given the increasing issue of antibiotic resistance.

The antibiotic ciprofloxacin, used as a control, showed substantial inhibitory effects on Lactobacillus spp. (34 mm zone of inhibition). This contrasts with most studies indicating Lactobacillus resistance to ciprofloxacin and suggests species-specific variability. The depletion of beneficial Lactobacillus spp. due to indiscriminate antibiotic use could lead to an increased risk of infections by opportunistic pathogens. This underscores the importance of cautious antibiotic use and the potential of natural alternatives like clove extracts to preserve the beneficial vaginal flora.

The MIC results further validated the antimicrobial efficacy of ethanol clove extract. Candida albicans showed the lowest MIC value (10 mg/ml), indicating higher sensitivity, while Staphylococcus aureus and Escherichia coli had MIC values of 20 mg/ml. These findings highlight the potential of clove extract as an effective antimicrobial agent against common vaginal pathogens, offering a promising alternative to conventional treatments.

The study successfully isolated and identified various bacterial and fungal species from vaginal swabs, demonstrating the microbial diversity within the vaginal environment. The significant antimicrobial activity of ethanol clove extract against key vaginal pathogens suggests its potential as an alternative therapy to antibiotics. This is particularly important in the context of increasing antibiotic resistance and the need for sustainable antimicrobial strategies. Future research should focus on optimizing the extraction methods and exploring the clinical applications of clove extract in managing vaginal infections. The findings emphasize the importance of maintaining a balanced vaginal microbiome and the potential role of natural products in preserving vaginal health.

